# *Drosophila* learn efficient paths to a food source

**DOI:** 10.1101/033969

**Authors:** Rapeechai Navawongse, Deepak Choudhury, Marlena Raczkowska, James Charles Stewart, Terrence Lim, Mashiur Rahman, Alicia Guek Geok Toh, Zhiping Wang, Adam Claridge-Chang

**Affiliations:** Institute of Molecular and Cell Biology, 61 Biopolis Drive, Singapore 138673; Singapore Institute of Manufacturing Technology, 71 Nanyang Drive, Singapore 638075; Duke-NUS Medical School, 61 Biopolis Drive, Singapore 138673; Department of Physiology, Yong Loo Lin School of Medicine, National University of Singapore, Singapore 138673; Sieva Pte Ltd

**Keywords:** *Drosophila*, flies, behavior, Skinner box, feeding task, learning assay, drug screening

## Abstract

Elucidating the genetic, and neuronal bases for learned behavior is a central problem in neuroscience. A leading system for neurogenetic discovery is the vinegar fly *Drosophila melanogaster*; fly memory research has identified genes and circuits that mediate aversive and appetitive learning. However, methods to study adaptive food-seeking behavior in this animal have lagged decades behind rodent feeding analysis, largely due to the challenges presented by their small scale. There is currently no method to dynamically control flies’ access to food. In rodents, protocols that use dynamic food delivery are a central element of experimental paradigms that date back to the influential work of Skinner. This method is still commonly used in the analysis of learning, memory, addiction, feeding, and many other subjects in experimental psychology. The difficulty of microscale food delivery means this is not a technique used in fly behavior. In the present manuscript we describe a microfluidic chip integrated with machine vision and automation to dynamically control defined liquid food presentations and sensory stimuli. Strikingly, repeated presentations of food at a fixed location produced improvements in path efficiency during food approach. This shows that improved path choice is a learned behavior. Active control of food availability using this microfluidic system is a valuable addition to the methods currently available for the analysis of learned feeding behavior in flies.

## Introduction

Learning and memory are fundamental brain functions that are important to all aspects of the human experience by allowing us to adapt to a challenging, changing environment. The molecular pathways underlying learning & memory are involved in both aspects of the ‘genes + environment’ sum, influencing our adaptability as individuals and serving as a conduit as we are shaped by experience. A better understanding of learning will be valuable to better treatment of addiction disorders and other forms of dysfunctional learning. Addiction disorders include food addiction, binge eating, and binge eating disorder (Marcus & Wildes, 2014), behavioral disorders that contribute to the worldwide obesity epidemic (Finkelstein & Strombotne, 2010). Obesity is a major risk factor for heart disease, stroke, type II diabetes, osteoarthritis, and some forms of cancer; public health policies have failed to reverse the epidemic and anti-obesity drugs have weak efficacy, problematic side effects, or both (Finkelstein & Strombotne, 2010; Gautron, Elmquist, & Williams, 2015). Finding better ways to treat obesity will require multidisciplinary efforts including basic research to connect dysfunctional reward learning with the neuroscience of hunger and satiety

An important model system for understanding the fundamental molecular and neural mechanisms of learning is *Drosophila melanogaster* (Keene & Waddell, 2007). Landmarks include the discovery of the first learning mutants (Dudai, Jan, Byers, Quinn, & Benzer, 1976), cloning of the associated genes, identification of the brain region that stores olfactory memories (Han, Levin, Reed, & Davis, 1992; Zars, Fischer, Schulz, & Heisenberg, 2000) and the circuitries that mediate aversive (Claridge-Chang et al., 2009) and appetitive (Burke et al., 2012; C. Liu et al., 2012) conditioning signals. In addition to memory research, *Drosophila* genetics has emerged as a powerful system to study other basic brain and metabolic functions, including food seeking (Sokolowski, 1980), food receptiveness (Deak, 1976), fat accumulation (Pospisilik et al., 2010), alcohol susceptibility (Moore et al., 1998), alcohol reward (Kaun, Azanchi, Maung, Hirsh, & Heberlein, 2011), and feeding regulation (Pool & Scott, 2014). Existing *Drosophila* assays have enabled major advances, but none currently give detailed information about behavior in response to single packets of food. There are a range of methods to measure various aspects of feeding behavior in larval and adult flies (Deshpande et al., 2014; Itskov et al., 2014; Ro, Harvanek, & Pletcher, 2014; Smith, Thomas, Liu, Li, & Moran, 2014), but none enable the automated control of food availability in freely moving adult flies. Active control of food availability is a long-established method in rodents (Skinner, 1930), but the adult male mouse weighs about 30 g while the adult male vinegar fly weighs about 0.6 mg, a 50,000-fold difference in size.

We developed a microfluidic feeder for *Drosophila* that delivers meal-sized, nanoliter-scale portions to a behavior chamber with visual and auditory stimuli. With repeated presentations, flies learned to approach food via more direct paths.

## Methods and Materials

### Drosophila

*Drosophila melanogaster* flies (a *yw* stock) were cultured in plastic vials at 22ºC, 60-70% relative humidity, under 12:12 hour light and dark cycles.

### Design of the SNAC microfluidic chip

Chips were designed with SolidWorks 2013 CAD software (Dassault Systemes, USA). The chip’s external dimensions were 33 mm × 30 mm × 4 mm; the behavior chamber was 20 mm × 15 mm × 2 mm (Figure 1A). The food channel delivered liquid to a feeding alcove; the volume of food delivered from this channel on actuation was 80 nL [range 60, 100]. We refer to the chip as the **S**mall-animal **N**utritional **A**ccess **C**ontrol (SNAC) chip. A fly’s head is ∼1 mm wide and its proboscis is <400 µm wide. The design aims were to control the liquid food delivery dynamically, allow video recording of both behavior and the microfluidic food channel. The design incorporated a feeding alcove that required that a fly insert its head to in order to drink from a feeding channel (Figure 1B). The channel was 200 µm wide and 50 µm deep, while the alcove was designed to be 400 µm wide. Completed chips showed a <20 µm divergence from the design dimensions for channel and ∼50 µm for the feeding alcove (Figure 1C).

**Figure 1.**
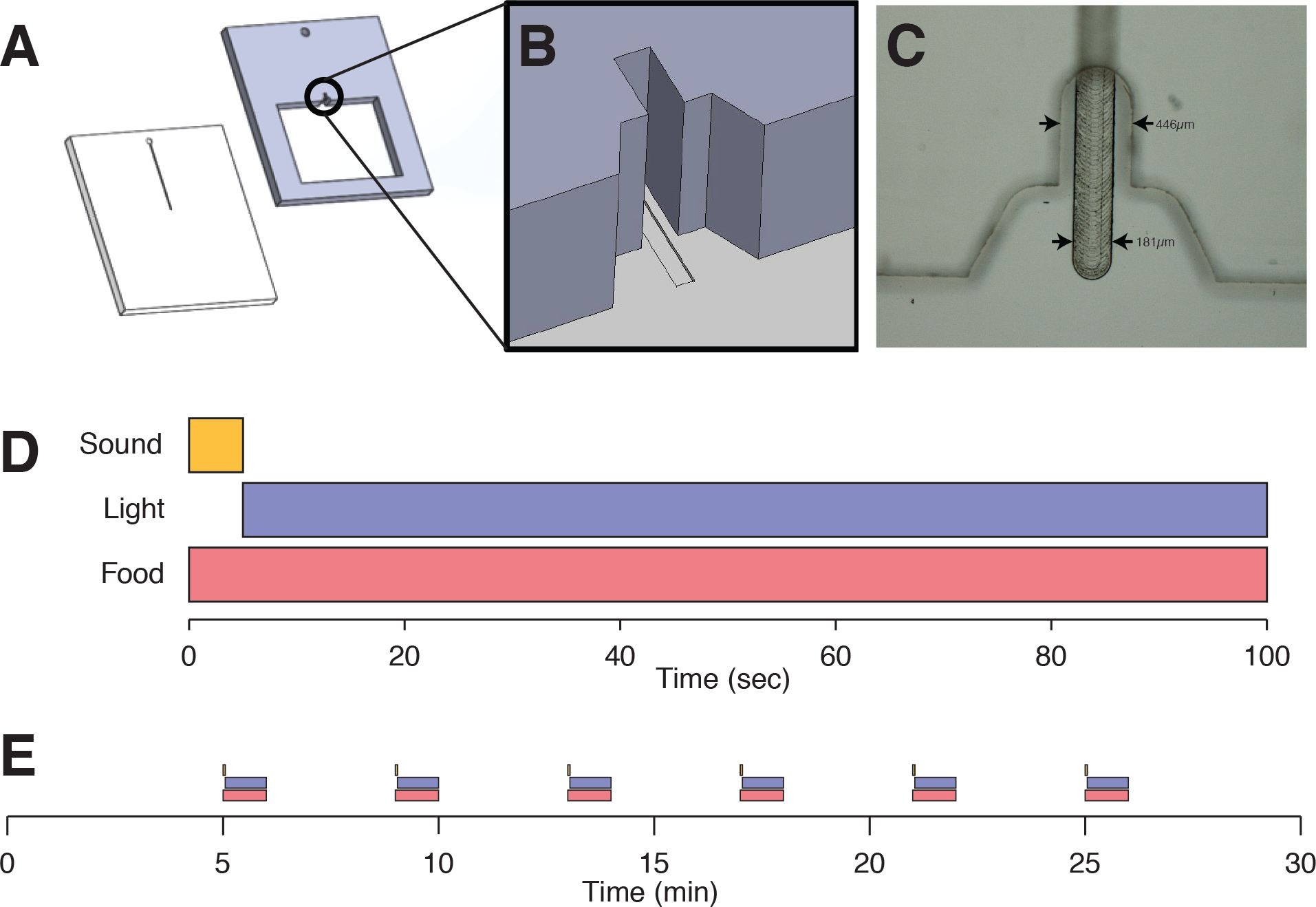
The Small-animal Nutritional Access Control (SNAC) microfluidic chip for food delivery experiments. A. The chip was milled from two pieces of polymethyl methacrylate (PMMA) before bonding. B. A view of the chip design showing the feeding alcove. The channel width was designed to be 200µm, the inner alcove was designed to be 400µm wide. C. A micrograph of the alcove and the food channel. The measurements after bonding were 446 ± 8µm alcove width and 181 ± 3 µm channel width (N = 5 chips). To drink the liquid, flies extended their proboscis into the narrow section of the alcove. D. A single food-availability epoch. It lasted up to 100 seconds of food delivery with a sound for 2 s and blue light that was kept on until the fly’s head was detected in the alcove. There was typically a 1-2s delay before food was extruded to be accessible. Control experiments omitted sound, light, or both; in sham trials, food was pumped close to the lick-port, visible but unreachable. E. Each group was subjected to six training epochs with intervening waits. Six feeding epochs each of 100 s duration were imposed, with 140 s wait intervals between the end of each epoch and the beginning of the next, a 4 minute epoch cycle.

### Chip fabrication and assembly

Chips were fabricated with optically clear thermoplastic cast polymethyl methacrylate (PMMA, Professional Plastics, Singapore). Computer numerical control machining was used to fabricate the chip layers. Valves and interconnects were made from polydimethylsiloxane (PDMS), cast from a pre-fabricated PMMA mold. The chip layers were bonded by thermal fusion as follows. The two chip layers (each 2 mm thick) were aligned in an L-shaped guide under an inverted microscope (Figure S1A). A small amount of acrylic glue was applied to the layer sides to hold them during thermal bonding (Figure S1B). The layers were sandwiched between 3 mm thick borosilicate glass sheets and tightened with binder clips (Figure S1C). This assembly was placed in a hot air bonding oven at 125ºC for 45 minutes, with a 1 h cooling time. The channel dimensions and bonding fidelity were measured with a 3D optical profiler (Zeta-20, Zeta Instruments). The chips were also tested with food dyes (Winner Brand, Thailand) for flow and leaks. Twenty chips were measured with the optical profiler at 40× objective to assess how closely they conformed with the design. After bonding, the alcove width was 446 ± 8 µm; the channel width and depth had the dimensions 181.4 ± 3.4 µm and 56.4 ± 6.3 µm, respectively. Error values are given as standard deviations.

**Figure S1.**
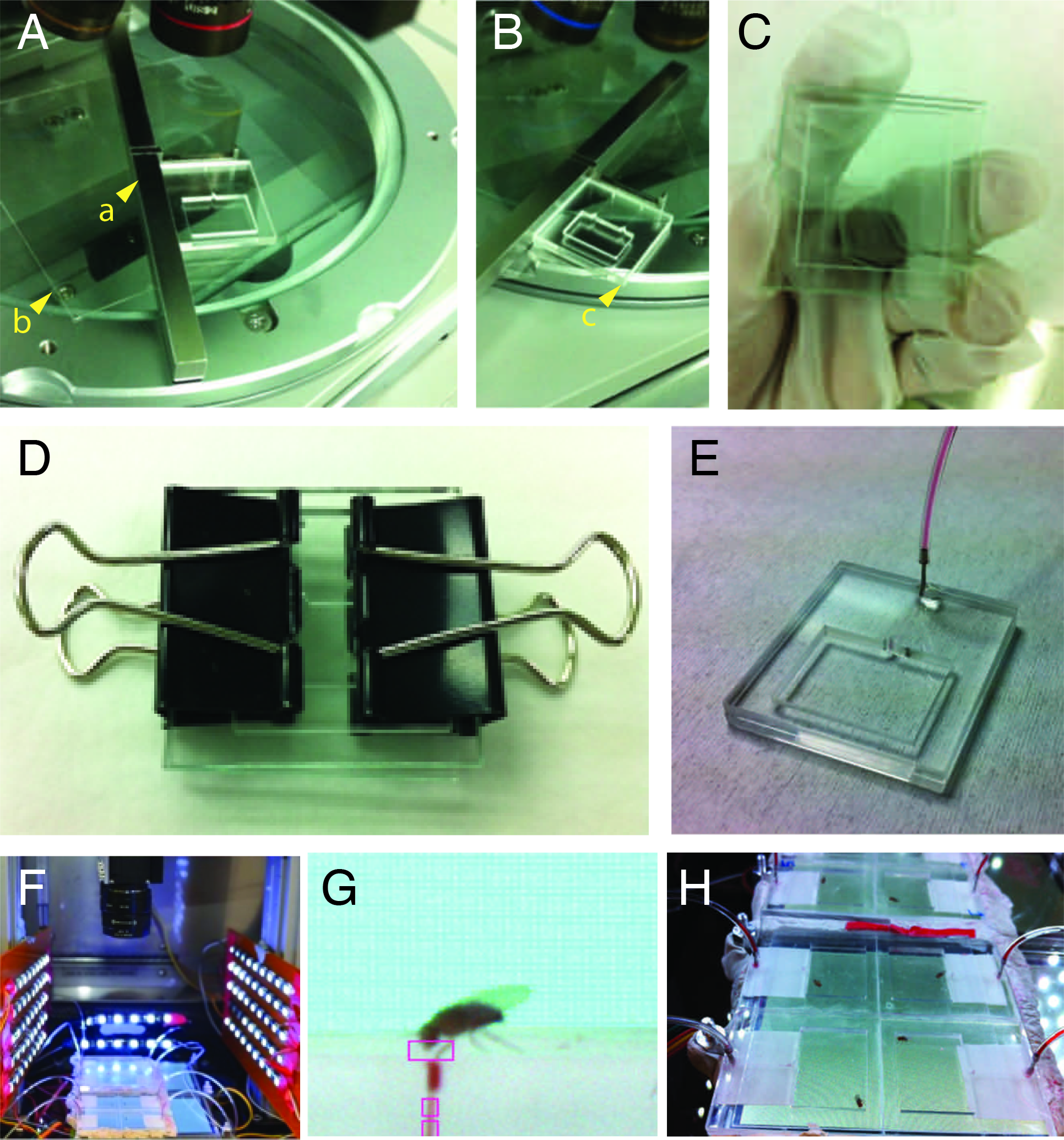
Fabrication, assembly and integration of the SNAC chip. A. Chip component layers were aligned with a metal L-guide (a) on top of a flat sheet of glass (b) under a microscope. B. Glue was applied to the corners (c) to maintain alignment during thermal bonding. C. Chip parts were sandwiched between two glass plates. D. Sandwich assembly was compressed with binder clips for oven bonding. E. The bonded chip was installed with a stainless steel tubing inlet. F. View of the system inside a temperature-controlled incubator. The cameras are in the top center of the image, directed at the chip 8-plex. LED arrays were positioned around the chips for illumination. The speaker was positioned directly below the chip-screen assembly. G. A fly at the feeding alcove of a single chip. Rectangles indicate regions of interest from the machine vision-pump control software: the top rectangle indicates the area of the feeding alcove being monitored for fly head presence, the two lower rectangles indicate control points for pump switching. H. A view of the SNAC chip 8-plex during operation. White tape was fixed to the bottom side of each chip for improved imaging contrast.

### Pumps and controllers

The SNAC chip system is shown in Figure S1F. A liquid food solution containing 5% sucrose (Sigma-Aldrich) and food dye (Winner Brand) was pushed and retracted through a microfluidic channel with custom syringe pumps (not shown). Each pump was constructed from a 10 µl precision glass syringe (80300, Hamilton, USA) to a 100mm linear actuator (L12 NXT, Firgelli, Canada). Each syringe was connected by flexible tubing (Tygon S-54-HL AAQ04103, OD 1/16”, Professional Plastics, Singapore) to a 4 mm long 21 G stainless steel tube inserted into the chip inlet. The tube was filled with mineral oil (Sigma M8410, U.S.) before adding sucrose solution from the chip-facing end. A microcontroller (NXP LPC1768, mbed, USA) was used to to drive the actuators with 5-volt digital pulses; pump speeds were adjusted by varying the pulse duty cycle. An H-driver circuit (SN754410, Texas Instruments, USA) was used to control pump direction. The microcontrollers were controlled with custom C++ firmware. During the experiment, fluid was alternately dispensed into the food channel and retracted back to a ’standby’ position. The fluid’s extent was detected by software that monitored color changes at distinct positions along the food channel (Figure S1G).

### System integration and sensory stimuli

Experiments were conducted with an eight-chip array (Figure S1H) in a temperature-controlled incubator (MIR-154, Sanyo, Japan). For light stimulus control, the chips were positioned on two LCD screens (µLCD-43, 4D systems, Australia) mounted on an aluminum stand. For sound stimuli, a 0.5W (8 ohm) speaker (COM-09151, Sparkfun.com) was mounted next to each screen. The chips were illuminated with white LED strips (ST-6500-CT, Inspired LED, U.S.A.) at 600 lux (measured on the chip surface). Two color cameras (A601fc, Basler, Germany) monitored animal activity and fluid location on all chips. All devices were controlled with a custom program in LabVIEW (National Instruments, USA).

### Experimental protocols

Four to seven day-old flies of both sexes were starved in batches of 10 for 24 hours in vials containing water-soaked tissue (Kimwipes). Flies were maintained in a 12:12 hour light and dark cycle at 22ºC during starvation. Flies were anesthetized on ice for less than a minute and transferred individually to chips. Two protocol variants were used. In Experiment 1, the flies were tracked 30 min before delivery and 60 min after delivery of a single food bolus. In Experiment 2, food was repeatedly delivered along with sound and light stimuli. Each 100s epoch contained a 2s 300Hz 82dB sound signal followed immediately by a white to blue screen change (Figure 1D). A food bolus of ∼80 nl (range 60-100) was delivered ∼3 s (range 1-5) after sensory cue onset. The screen was kept blue until the fly’s head was detected to be in the feeding alcove, upon which the food was retracted and the screen returned to white. Six food/stimulus epochs were presented over 30 minutes (Figure 1E) in a 4 minute cycle, with a 140 s wait between epochs.

### Tracking, feeding metrics and data analysis

The animals were tracked with computer vision code in LabVIEW using standard background subtraction and centroid methods. For tracking in changing blue/white light conditions, the red or blue plane was extracted from the color video when the screen was white or blue, respectively. The alcove width in each video was used to rescale tracking data to millimeters. Behavior data were plotted with Matlab; summary statistics were means or median with relevant confidence intervals shown as error bars. ‘Time to alcove’ measured the time the animal spent after food presentation and before detection of a fly’s head in the food alcove. The path efficiency was calculated as the distance of the most direct path to the feeding alcove divided by the actual distance travelled by the fly during a feeding epoch, a measure of how directly flies moved to the food from their location at the start of an epoch, with a figure closer to 1 indicating a more direct path. Time to alcove and path efficiency were only computed on trials in which an alcove head detection event occurred. One-Way ANOVA tests were used for comparisons over epochs. Estimates are reported as means with their 95% confidence interval (‘95CI’) following the convention [lower bound, upper bound]. Both raw mean difference effect sizes and standardized effect sizes (Hedges’ *g*) were used to estimate the magnitude of effects. Hedges’ *g* estimates the change between two groups in terms of their standard deviations, i.e. *g* = 1 indicates a one standard deviation shift between groups. T-tests were used to calculate p values for comparisons of two independent groups.

## Results

### Flies briefly increased their locomotion around food intake

We examined behavior before and after consumption of a single bolus of a 5% sucrose solution. An example of the behavior observed is illustrated in Figure 2A and in Supplementary Video 1. Baseline median walking speed was less than 0.1 mm/s, but increased sharply when the food bolus was discovered by the fly and walking speed remained high for several minutes after feeding (Figure 2C,D). We conclude that flies undergo locomotor arousal around a feed event.

**Figure 2.**
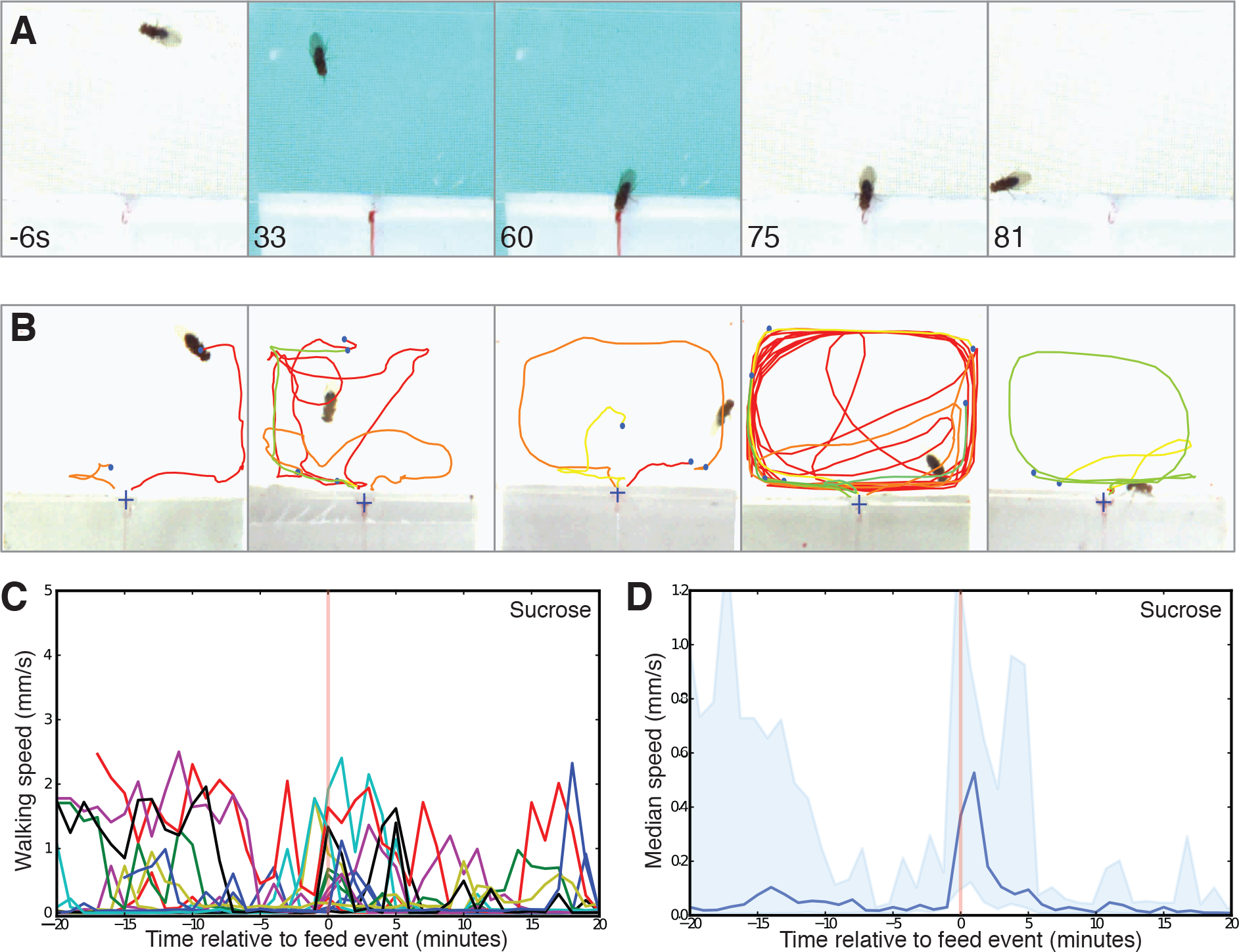
Fly walking activity responds to a food bolus. A. Five video frames of a fly in a single food presentation. Before the food delivery, the fly is walking around a white-illuminated chamber (-6 s). Blue light is activated and food is extruded, the fly has not approached the alcove at 33 s. At 60 s the fly enters the alcove and feeds, which triggers the software to retract the food and switch the screen back to white. The fly stays eating residual food before leaving and returning to walking around the chamber (81 s). B. Cumulative traces of five flies; blue dots indicate the location of the fly at the start of the epoch, ‘+’ symbols indicate the alcove location. Each coloured trace represents the path taken during one epoch period and only epochs where the fly entered the feed alcove are shown. C. Individual fly walking speeds before and after feeding on a bolus of sucrose liquid food. Tracking data timelines were re-centered around feeding events. D. Median walking speed of flies fed sucrose food. Light blue error band indicates confidence intervals of the median. Walking speed is affected by food intake. The pale vertical red line indicates the time of feeding events.

### Flies were able to discriminate accessible from inaccessible food

We aimed to identify learned aspects of food approach after repeated presentations. Starved flies were subjected to a six-epoch regime with food, a screen color change and a 2 second audio cue (Figure 1E). In this regime, when food was made accessible in the alcove, flies entered it an average of 3.6 out of a possible 6 epochs [95CI 3.3, 3.9] (Figure S2A). To investigate the cues that flies used in making a food approach, liquid food was pumped along the channel but stopped just before the food port. The latency to alcove entry (time to alcove) in each epoch was measured by detecting the presence of the animal’s head in the food port. When food was visible but inaccessible, flies entered the feeding alcove in an average of only 0.5 of 6 epochs [95CI 0.3, 0.7] (Figure S2A). Of the flies that made it to the alcove, there was little difference in behavior: the time to alcove for accessible food was 30.5 seconds [95CI 27.9, 33.1] and 39.1 seconds [95CI 28.4, 49.8] for inaccessible food (Hedges’ *g* = 0.34, p = 0.10; Figure S2B). In each epoch, a fly may be near or far from the food alcove, and in each case the direct path to the alcove is a straight line. The path efficiency was also similar for both conditions (Figure S2C). These results indicate that flies could usually discriminate inaccessible food from accessible, but that in the minority of cases where they approach the alcove, they do so with similar speed and efficiency.

### Background color changes and an auditory signal increased food approach

In conditioning chambers for vertebrate experiments, some protocols use one or more sensory stimulus to cue reward delivery. We asked whether flies were using the screen color change and the audio signal as cues for the alcove approach. Four experiments omitting either the color switch, the audio pulse, or both were performed: both blue light and a 300 Hz tone, light-only, sound-only, or neither stimulus. Path efficiency and time-to-alcove were largely unchanged by the presence of sensory cues (Figure S2E-G). When both stimuli were presented together, the number of alcove entries per fly was higher by at least one entry relative to either single-stimuli or no-stimulus conditions, (Figure S2D) (ANOVA p = 1.6 × 10^−7^). Thus, only the visual + auditory stimuli combination promoted increased food approach.

**Figure S2.**
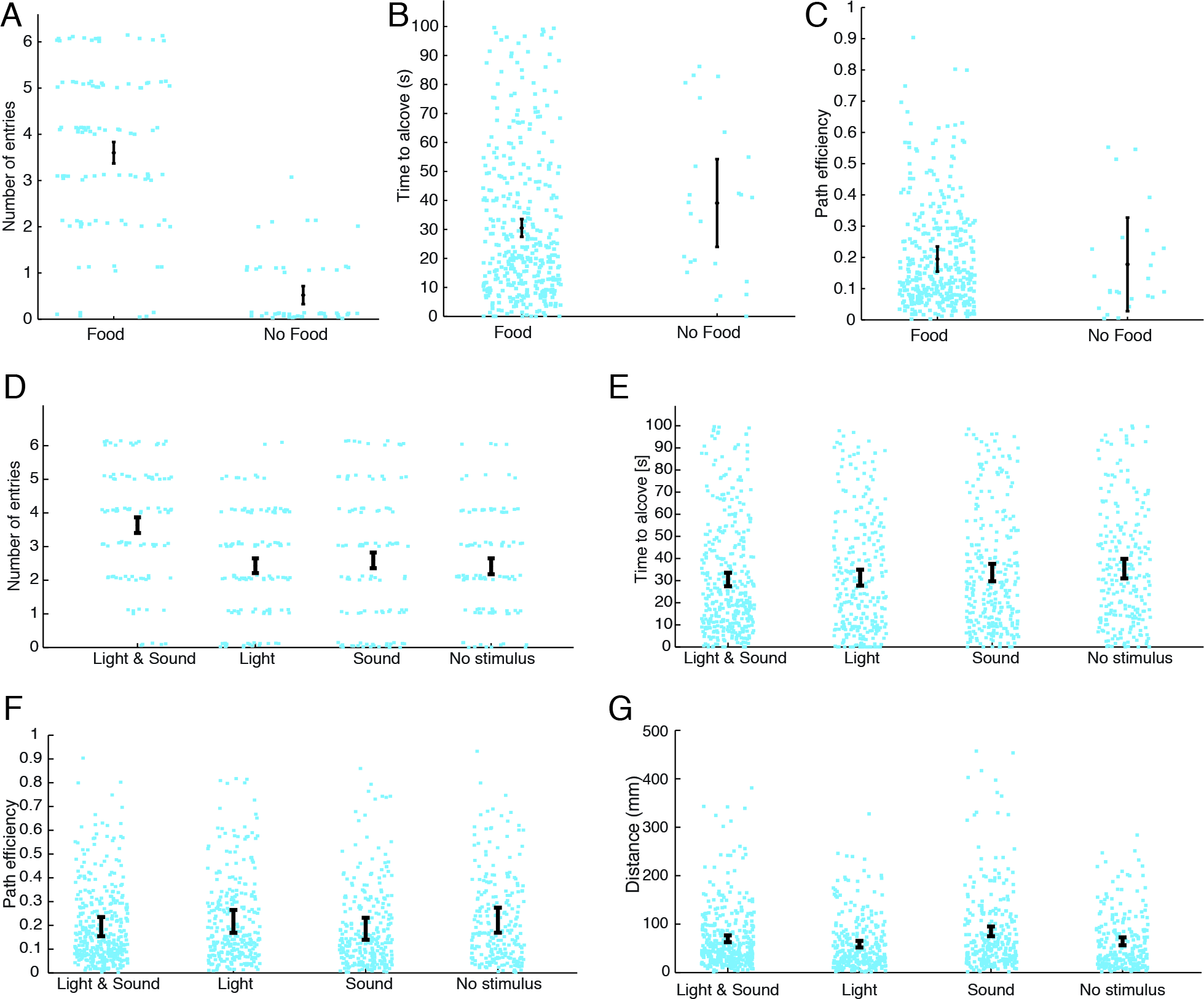
The effects of food delivery and sensory cues on feeding alcove approach. A. Flies enter the alcove more frequently when food is accessible. A comparison of experiments in which liquid food was extruded into the trough (N = 103) and experiments where the fluid was held inaccessible (N = 48) shows that visibility of food alone is poor at encouraging alcove entries. There was a large difference in the number of alcove entries (Hedges’ g = 2.1, contrast p < 0.0001). B. Whether food was accessible or only visible made little difference to the time to alcove entry for flies that did enter: a difference of only 8.6 seconds was observed (contrast p = 0.1). C. The path efficiency was similar in both food-accessible and food-inaccessible conditions: 0.19 path efficiency [95CI 0.18, 0.21] and 0.18 path efficiency [95CI 0.11, 0.24] respectively (contrast p = 0.61). D. Omitting one or both stimuli had moderate effects on alcove entries relative to the combination of light and sound; each stimulus alone had little or no effect relative to food presentation alone. Flies entered in 3.6/6 epochs [95CI 3.3, 4.0] with both stimuli versus 2.4/6 [95CI 2.1, 2.7] with light-only, 2.6/6 [95CI 2.2, 2.97] with sound-only, and 2.4 of 6 [95CI 2.1, 2.7] with no stimuli. E. The time to alcove was largely unaffected by the presence of stimuli: flies given both cues took an average 30.51 s [95CI 27.9, 33.1] to enter the alcove (N = 371 epochs). Flies in other conditions took only slightly longer; light only 31.3 s [95CI 28.2, 34.4] (N = 282), sound only 33.7 s [95CI 30.5, 36.8] (N = 272), no stimuli 35.4 s [95CI 32.0, 38.7] (N = 239). Time to alcove scores were only counted in epochs where a fly entered the alcove. F. Path efficiency was 0.19 [95CI 0.17, 0.21] when both light and sound were used to cue the epoch start, and this was no different from a blue light-only cue 0.22 [95CI 0.2, 0.24], audio stimulus 0.19 [95CI 0.17, 0.21] and no sensory cues 0.22 [95CI 0.2, 0.24]. G. The total distance travelled in a 100-s epoch varied only slightly between all conditions. Flies moved the least in the light cue only condition (58.7 mm [95CI 52.3, 65.0]; Figure 6D), and the furthest in the sound cue only condition (85.1 mm [95CI 75.0, 95.2]; Figure 6D). Surprisingly, the reference light and sound condition did not differ substantially from no stimuli (69.8 mm [95CI 63.3, 76.2] and 64.6 mm [95CI 55.0, 74.1], respectively).

### Over repeated presentations, food approach and time to food were unchanged

Two behavioral metrics, the proportion of flies entering at the alcove and the time to the food alcove, were analyzed over epoch number (Figure 3A-B). Only modest, non-statistically significant differences in the proportion of alcove entries were observed (Figure 3A). A modest dip in the time to alcove was observed by the third epoch, but there was no statistical change in this metric by the sixth epoch (Figure 3B). These results indicate that the flies’ frequency of food approach and the time taken to approach a food source undergo little or no adaptation during repeated presentations. Both male and female flies were tested in these experiments, no substantial differences in the proportion of alcove entries were observed between these groups (data not shown).

**Figure 3.**
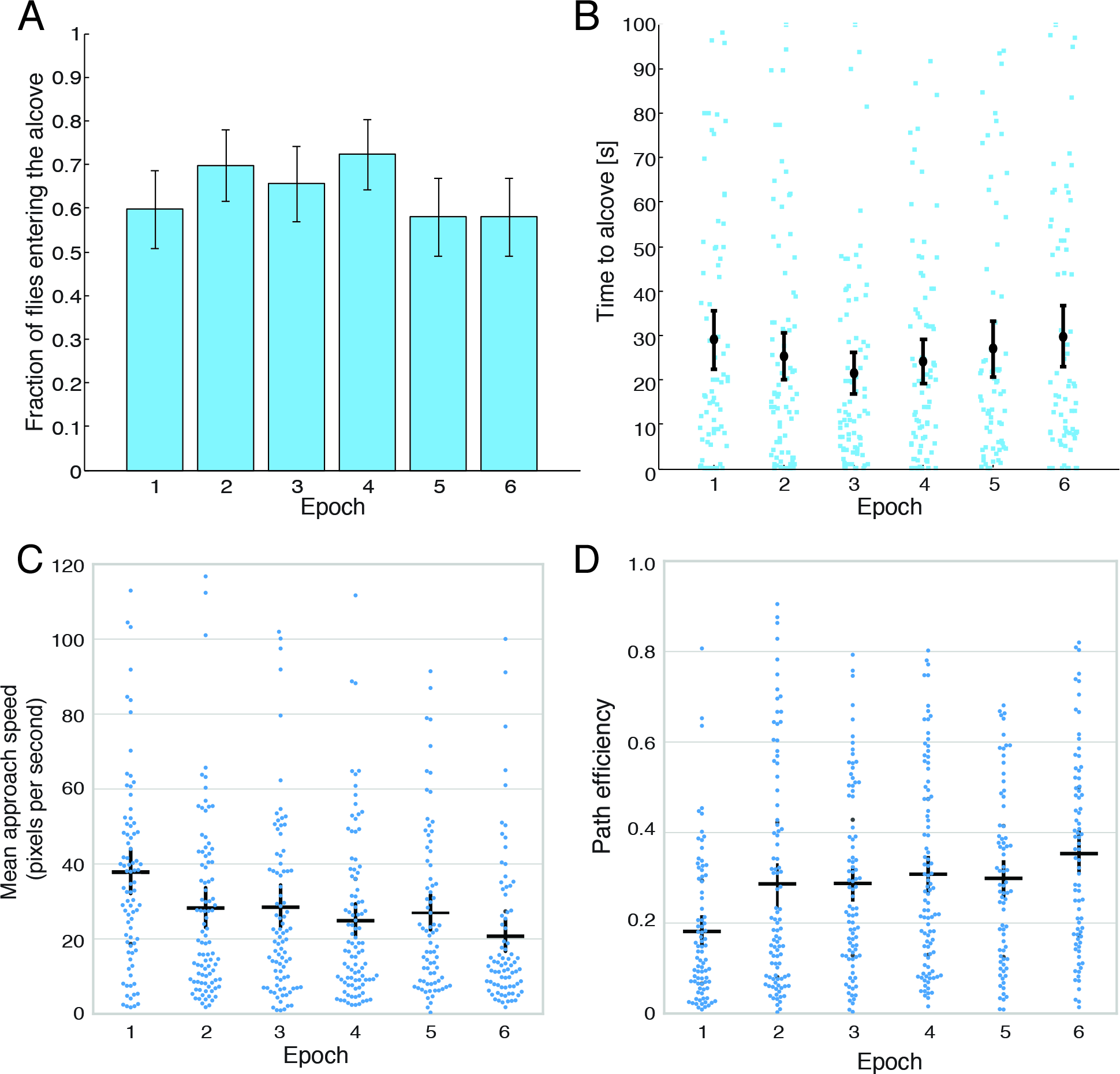
Feeding behavior of flies over six epochs of food presentation. A. The proportion of flies that entered the alcove did not change in a consistent direction over six epochs. B. The time latency to approach the alcove for flies in each of six 100-s epochs decreased until the third epoch before returning to the original time. C. Mean alcove approach speed decreased over six epochs. D. Path efficiency increased over six epochs by 1.06 *g*, a large effect.

### Flies learn to improve their food approach path efficiency

Flies generally did not follow direct paths to the food after the epoch commenced, but displayed more or less circuitous paths during each food presentation epoch (Figure 2B). Flies’ alcove approach path efficiency increased progressively over repeated presentations, from 0.18 in the first epoch to 0.34 by the sixth epoch, a 0.16 path efficiency increase [95CI 0.10, 0.22] (Figure 3D). The standardized effect size indicated that the flies made a large improvement in path efficiency (Hedges’ *g* = 1.06, p < 0.0001). These data indicate that flies learn to follow more direct paths to a food delivery location over repeated presentations.

### Path efficiency is weakly related to walking speed

Path efficiency increases despite a largely unchanged time take to enter the alcove (Figure 3B, suggesting that the flies are moving more slowly towards the alcove in more efficient epochs. A plot of mean alcove approach speed over epoch confirmed that flies moved slower in later, more efficient epochs (Figure 3C). To investigate how closely path efficiency was related to walking speed, we performed a linear regression of the two metrics, finding that there was indeed a relationship, albeit a weak one, R^2^ = 0.12 (Figure S3A). We also found that path efficiency was related to path length, R^2^ = 0.27 (Figure S3B). Thus, in successive food presentation epochs, flies tend to walk shorter paths towards the alcove more efficiently and more slowly.

**Figure S3.**
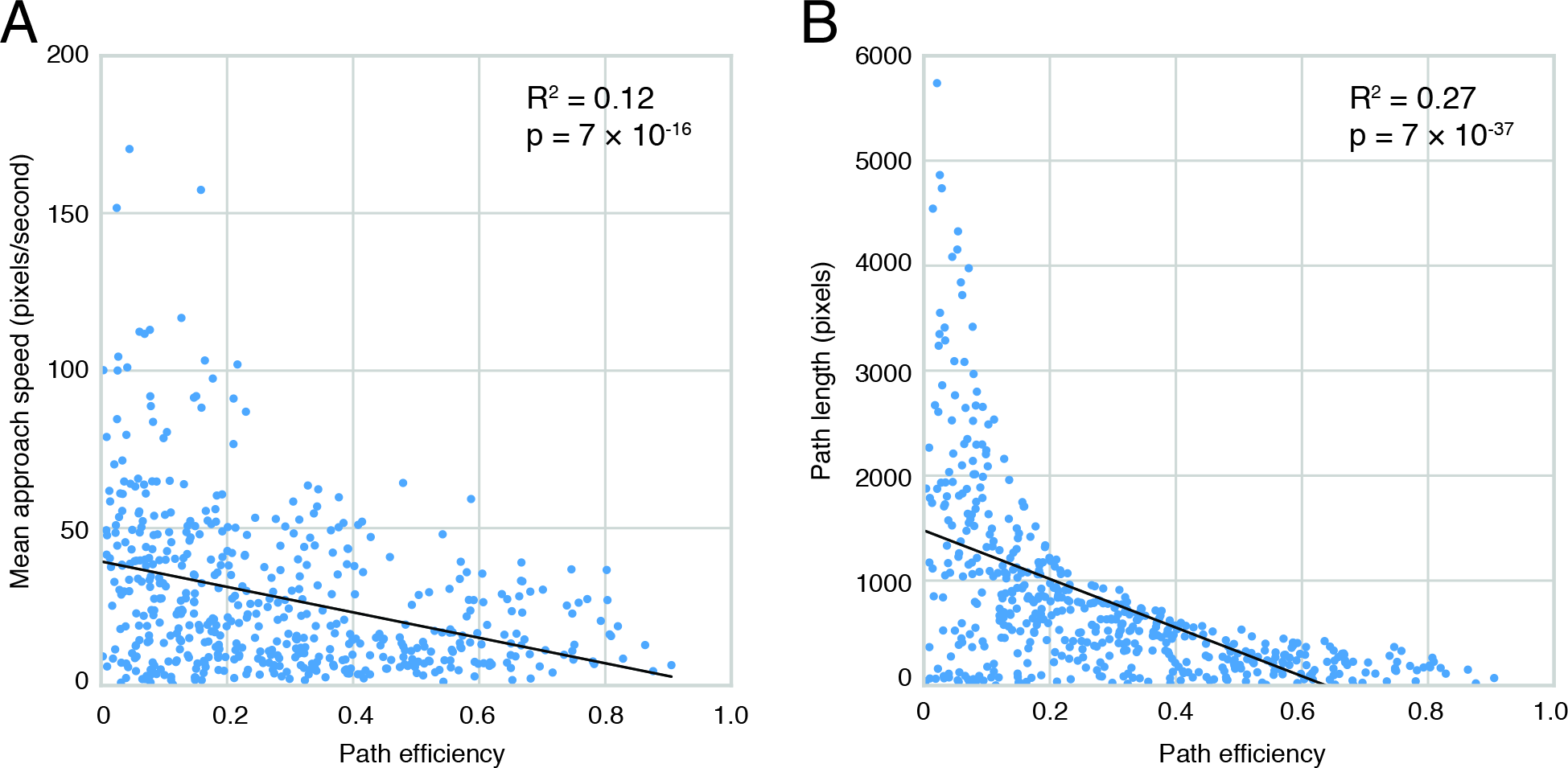
Relationships between path efficiency, speed and path length. **A.** Path efficiency and mean alcove approach speed are weakly correlated, R^2^ = 0.12. **B.** Path efficiency and path length to the alcove are correlated, R^2^ = 0.27.

## Discussion

The tiny size of the genetically tractable insect *Drosophila melanogaster* means that some tools development has lagged behind some available for rodent species. A number of innovative methods to study fly feeding are currently available, including CAFE (Ja et al., 2007), which uses food capillaries to provide a precise quantification of food intake, and flyPAD (Itskov et al., 2014) and FLIC (Ro et al., 2014), which use electrical methods to detect food contact events with high temporal precision, but there are no methods to dynamically control access to defined quantities of food in fly. Here we show that the SNAC microfluidic chip enables the delivery of small quantities of liquid food (∼80 nanoliters) to flies while simultaneously tracking animal locomotion, allowing the system to capture animal behavior when food is presented. The utility of the SNAC chip system can be further enhanced by the addition of components that enable computer vision feedback to control food access in response to the animal’s behavior, in a similar manner to a ‘Skinner’ conditioning apparatus.

While we found no evidence for behavioral adaptation for the fraction of flies entering the alcove to feed or the time taken to reach the alcove, we found that flies learn to walk along more efficient paths to a transient food source. Surprisingly, there is no relationship between path efficiency and time-to-alcove, and flies walk more slowly towards the alcove in later epochs. These results indicate that, on average, flies slowly follow more direct paths to the feeder during later epochs, rather than the rapid exploration that occurs in the earlier epochs. That they learn to take more efficient paths shows that flies associate food with a location within an enclosed space. This result is compatible with results showing that flies can associate food with odors (Krashes & Waddell, 2008), and are capable of visual place learning (Ofstad, Zuker, & Reiser, 2011). Previous studies on larval foraging behavior showed that genetic functions are shared between foraging and learning (Mery, Belay, So, Sokolowski, & Kawecki, 2007). Path efficiency learning may be relevant to foraging adaptation in wild *Drosophila* adults. The development of a microfluidic dynamic feeder device for feeding and learning analysis opens new possibilities in the study of learned foraging behaviors.

## Acknowledgments

This work received support from A*STAR Joint Council Office Grant 1231AFG030 awarded to ACC at IMCB and ZPW at SIMTech. ACC, RN, JCS, M. Raczkowska and M. Rahman were supported by a Biomedical Research Council block grant to the Neuroscience Research Partnership and IMCB. ZPW, DC, TL, and AGGT were supported by a Science and Engineering Research Council block grant to SIMTech. ACC received support from Duke-NUS Medical School. We thank Sharon Tan Khai Hoon and Rahmad Bin Selamat for assistance with chip fabrication, Anders Eriksson for manuscript comments and Mai Adachi for data analysis help. We thank Kerry McLaughlin of Insight Editing London for critical review of the manuscript.

